# A pancreas specific *Ptf1a*-driven Cre mouse line causes paternally transmitted germline recombination

**DOI:** 10.1101/2020.03.13.989178

**Authors:** Derya Kabacaoglu, Marina Lesina, Hana Algül

**Author notes:** To whom correspondence should be addressed: Hana Algül, Klinikum rechts der Isar, Technische Universität München, Ismaninger Str. 22, 81675 Munich, Germany.

## Abstract

Cre-*lox*P recombination system is a commonly used tool to achieve site-specific genetic manipulation in genome. For multiple Cre driver mouse lines, parental transmissions of recombined flox alleles are reported. *Ptf1a*-driven Cre lines are widely used to achieve genetic manipulation in a pancreas specific manner. Herein, we report germline recombination in breedings when *Cre* allele is retained paternally in *Ptf1a^tm1(cre)Hnak^*. The germline recombination frequency changed depending on the target allele. Therefore, unless the reporter allele is on the target gene, the reporter activity is to be validated. Overall, we highlight that all *Ptf1a*-driven Cre mouse lines should be genotyped for possible germline recombination and we advise the maternal transmission of the *Cre* to progeny.

## Introduction

Identified firstly in 1981, “Causing recombination” (Cre) is isolated from P1 bacteriophage, which is shown to recombine “*locus of crossing over (x1),* P1*”* (loxp) sites [1]. Since its discovery, use of Cre-*lox*P system is highly utilized in mouse models. Depending on the approach, it can be used to provide precision in both cell type and temporal control of the genetic manipulation.

Although the method is powerful, recent evidence suggests certain limitations. Cre-mediated cytotoxicity is reported for various cell types both *in vitro* and *in vivo* [2–11]. In these studies, postulated reasons are DNA damage induced by Cre-mediated genome-wide non-specific recombination events and altered cell signaling pathways (e.g. PKA) [12]. Additionally, organ-specific Cre expression is highly problematic due to limitations in identification of individual proteins getting expressed in specific cell types.

Multiple Cre lines have been developed in order to achieve pancreas-specific recombination in mouse models. Among many, one of the most used models is the line in which *Cre* is knocked in *Pancreas associated transcription factor 1a* (*Ptf1a*, a.k.a. *p48*) locus [13]. *Ptf1a* has an essential role during pancreatic epithelium generation. Further differentiation of sub-pancreatic compartments like acini, endocrine, and ducts are also regulated by *Ptf1a* [14–19].

In order to study *Ptf1a* function in the developing mouse, Nakhai et al. generated a model in which *Cre*-coding sequence was knocked in the Exon-1 of *Ptf1a* via homologous recombination (*Ptf1a^tm1(cre)Hnak^*). Several studies showed PTF1A expression in developing pancreas [16], neural tube [20], and cerebellum [21,22]. With the use of *Ptf1a^tm1(cre)Hnak^*mouse model, PTF1A was shown to be expressed also in the mouse retina in addition to the previously identified locations [13]. Interestingly, a recent study proposed a regulatory function of *Ptf1a* in hypothalamus for sexual development during embryonic stage [23]. Although there are many studies for the function of *Ptf1a* in the developing mouse, the ones focusing specifically on the reproductive system of adult mice are in scarce.

Here, we provide evidence for paternally-driven germline recombination mediated by Cre expression in *Ptf1a^tm1(cre)Hnak^*mouse line. The recombination event was observed in multiple floxed alleles, while the frequency varied depending on the target gene. Additionally, *Cre* allele transmission into progeny was not necessary for the germline recombination. Recent other reports also support parental transmission of germline recombination in many different Cre driver lines [24–34]. Therefore, precaution needs to be considered for the experimental design of the mouse studies. We propose that the *Cre* transmission should be performed only through females in which we observed no germline recombination in the progeny.

## Results

In order to study the function of *RelB* in pancreatic cancer, we aimed to generate a mouse model, which expresses KRAS^G12D^ and knocks out *RelB* in the pancreas. Therefore, using mouse models *Relb^tm1Fwei^*, *Ptf1a^tm1(cre)Hnak^*, and *Kras^tm4Tyj^* we followed a breeding scheme as provided in Figure 1A. Our genotyping primers for *RelB* were already designed to recognize all wt, flox, and recombined alleles (kindly provided by Marc Riemann), which made it possible for us to track-down the recombination in every pup. Initially, in F0 generation a *Relb^fl/fl^* male was put into breeding with 2 separate females either with a *Ptf1a-Cre/+* or *Kras^fl/+^*. In the F1 generation, females with either *Relb^fl/+^*; *Ptf1a-Cre/+* or *Relb^fl/+t^*;*Kras^fl/+^* were put into breeding with *Relb^fl/fl^* males in order to obtain homozygous *Relb^fl/fl^* allele with either *Ptf1a-Cre* or *Kras*. None of the progeny from these breedings had a recombined *Relb^fl^* allele (data not shown). And finally, in the F2 generation, we put a *Relb^fl/fl^*; *Ptf1a-Cre/+* male into breeding with a *Relb^fl/fl^*; *Kras^fl/+^* female. Interestingly, 28 of 28 pups had one recombined and one unrecombined flox alleles observed upon ear-genotyping (Figure 1B). The absence of germline recombination in F2 generation was validated with post-genotyping (Figure 1C). The germline recombination occurrence was irrespective of their *Cre* status (Figure 1B). This implies *Cre* transmission is not required for germline recombination in the descendants. To validate the whole-body recombination, mice with heterozygous germline recombination were sacrificed and their various organs were harvested for DNA extraction and PCR genotyping. The results show that all of the organs contained one recombined and one unrecombined *Relb* allele, while no *Kras* germline recombination was observed in the respective pups (Figure 1D).

**Figure 1:**
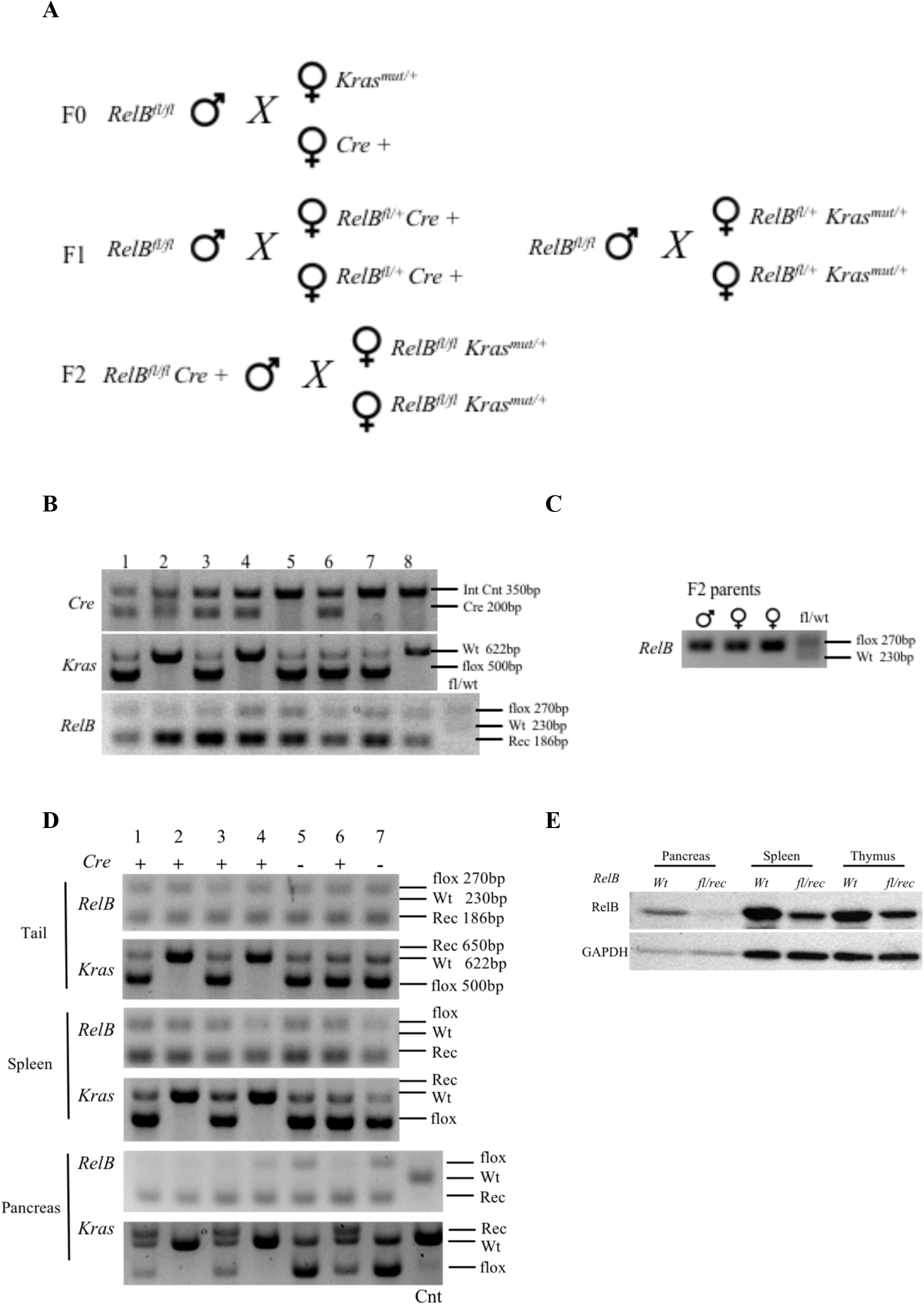
Germline recombination observed in *RelB^fl^* mice. Names of the primer pairs are indicated on the left. Lengths of the respective alleles are indicated on the right. A) Breeding scheme to obtain *Relb^fl/f^*; *Ptf1a-Cre/+*; *Kras^fl/+^* mice. B) Representative Genotyping PCR results of the pups obtained from F2 generation. C) Genotypes of the mice which were in F2 breeding. D) PCR results from tail, spleen and pancreas recognizing both the recombined and unrecombined alleles. D) RelB western blot results of thymus, spleen and pancreas in *RelB* germline recombined and *wt* tissues. **Rec**: recombined floxed allele. **Int Cnt**: internal control

The genotyping results of the pancreatic DNA implies a difference in the recombination efficiency in different floxed genes. While the recombination efficiency of the single-unrecombined floxed *RelB* allele is quite strong, *Kras* recombination is not that efficient. Immunoblot analysis of the various organs also showed a reduction in RelB expression (Figure 1E). However, since none of the alleles had a reporter activity, we couldn’t test additional measurements.

To see if the same recombination event occurred in different alleles we used two different mouse lines: *Rel^tm1Ukl^* and *GFP-Rel. Rel^tm1Ukl^* is a knock-in mouse model in which endogenous *Rel* Exon-1 locus is flanked by 2 *loxp* sequences. Upon recombination, exon 1 is recombined and a promoterless *GFP* gene meets with a *PGK* promoter. This way, *Rel* recombination can be efficiently traced down by GFP expression. GFP-Rel is a transgenic mouse model (kindly provided by Marc Schmidt-Supprian) in which a BAC construct with a *PGK* promoter controlled, N’ *GFP* tagged *Rel* is integrated into genome. Simplified maps along with the locations of the primers recognizing the recombined and unrecombined floxed alleles are shown in Supplementary Figure 1.

Breedings with *Rel^fl^* (*Rel^tm1Ukl^*) and *Ptf1a-Cre* carrying males indicated similar results as in *RelB* mice. In a retrospective PCR analysis of 3 different breeding pairs with *Rel^fl^*; *Ptf1a-Cre/+* fathers, 36/45 pups (80%) had recombined *Rel* allele, while 9 of them (20%) retained their unrecombined floxed *Rel* (for pups where 1 *Rel*^fl^ but not *wt* allele is paternally transferred) (Table 1). PCR from the different organs collected from various recombined mice also supported the presence of germline recombination (Figure 2A). Immunoblot analysis from the heterozygous recombined mice indicated eGFP expression only in the germline recombined mice and *Ptf1a-Cre/+* pancreas along with a reduced c-Rel expression (Figure 2B). Of note, breeding pairs with 6 different females with *Rel^tm1Ukl^*; *Ptf1a-Cre/+* showed no recombination in their progeny (0/93 pups, 0%) (data not shown).

**Table 1:**
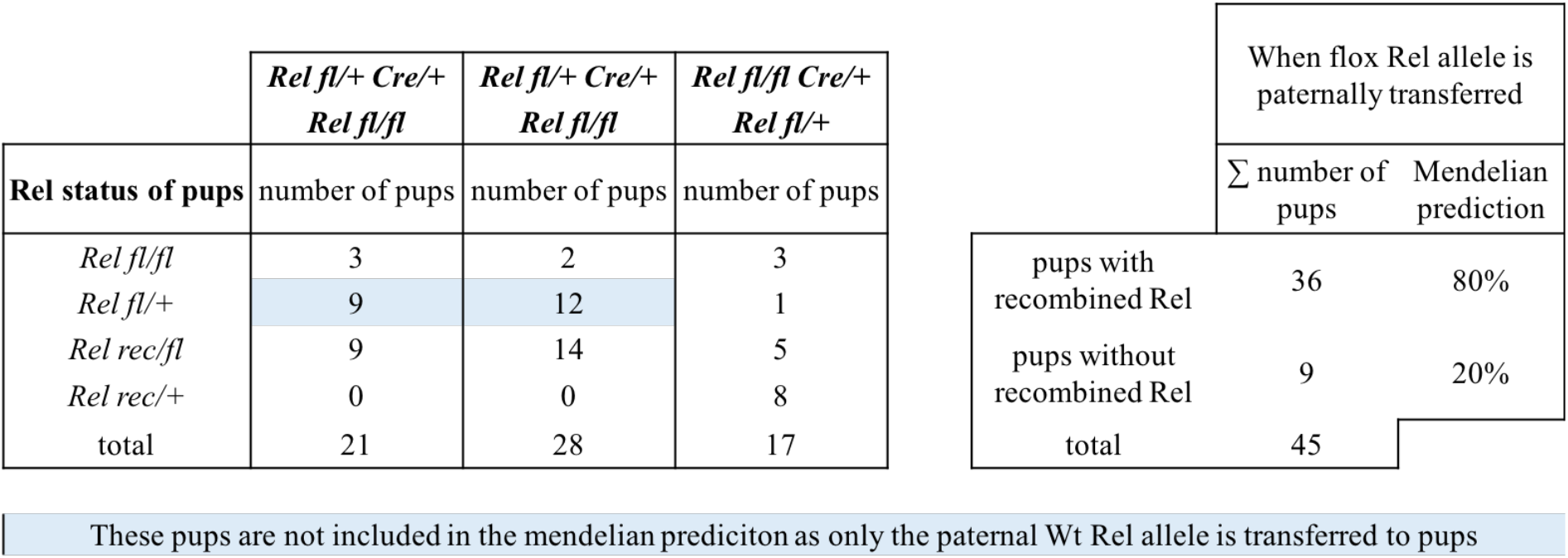
Number of mice obtained from *Rel* breeding

**Figure 2:**
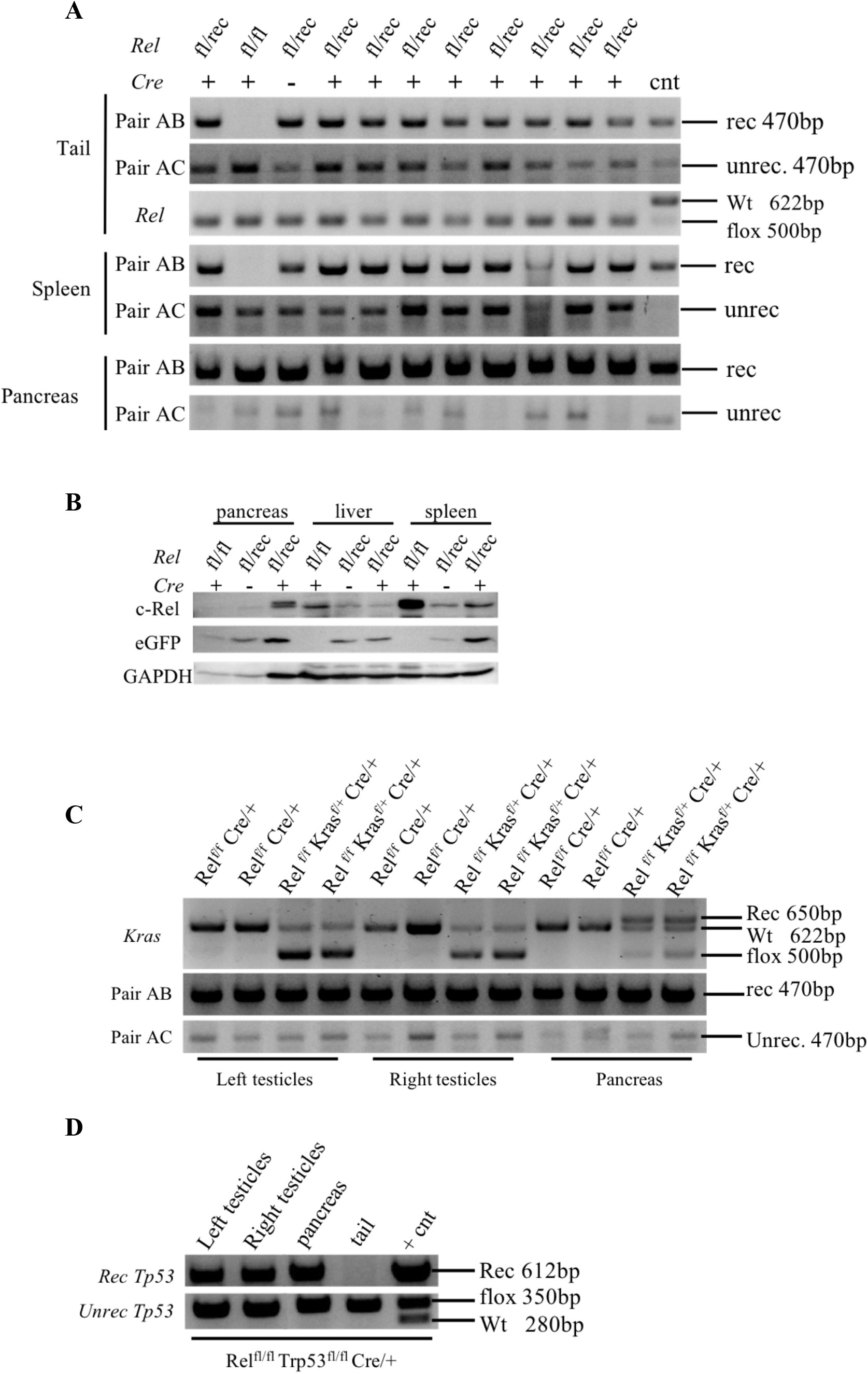
Germline recombination observed in *Rel^fl^* mice. A) Primer pairs A+B recognize the recombined allele, while the A+C recognizes the unrecombined floxed allele. *Rel* is the genotyping primer mix recognizing either floxed or wt alleles which can’t differentiate recombination. B) eGFP and c-Rel western blot results of pancreas, liver and spleen in *Rel* germline recombined and unrecombined tissues. C) PCR results for *Kras* and *Rel* recombination in testes and pancreas. D) PCR results for Tp53 recombination in testes pancreas and tail. **Rec**: recombined floxed allele. **Unrec**: unrecombined floxed allele.

In order to analyze if there was recombination in reproductive system of the males which were not germ-line recombined, we isolated DNA from testes and pancreas. PCR analysis showed recombination of *Rel* in both of the testes along with pancreas, although none of the mice had germline recombination. Interestingly, despite *Rel* was recombined, *Kras* was not (Figure 2C). *Tp53* is a very commonly used gene in mouse models to study pancreatic cancer. In order to test possible *Tp53* germline recombination in *Trp53^tm1Brn^* model, we genotyped 5 different breeding pairs in which Cre was paternally transmitted. We observed no *Tp53* germline recombination in these mice (data not shown). Interestingly, the male reproductive system still showed recombined *Tp53* allele (Figure 2D).

To eliminate the possibility of mosaic Cre recombination in the tail and ear biopsies, we took advantage of the *GFP-Rel* mouse model. Because it is a transgenic allele, it has been kept always in a heterozygous state, and always bred with a wildtype mate. To test possible mosaic recombination, a male with *GFP-Rel^tg/+^; Ptf1a-Cre/+* was bred with *Kras^fl/+^* females. A new set of primers were designed to recognize both unrecombined and recombined alleles (Supplementary Figure 1). Among 40 pups, 22 of them have *GFP-Rel* allele. 14 of 22 were regenotyped for *GFP-Rel* germ line recombination. 6/14 mice had *GFP-Rel* recombination in tail PCRs irrespective of *Cre* transmission, and there was no unrecombined *GFP-Rel* allele in these mice (Table 2) (Figure 3). This eliminates the possibility of a mosaic recombination.

**Table 2:**
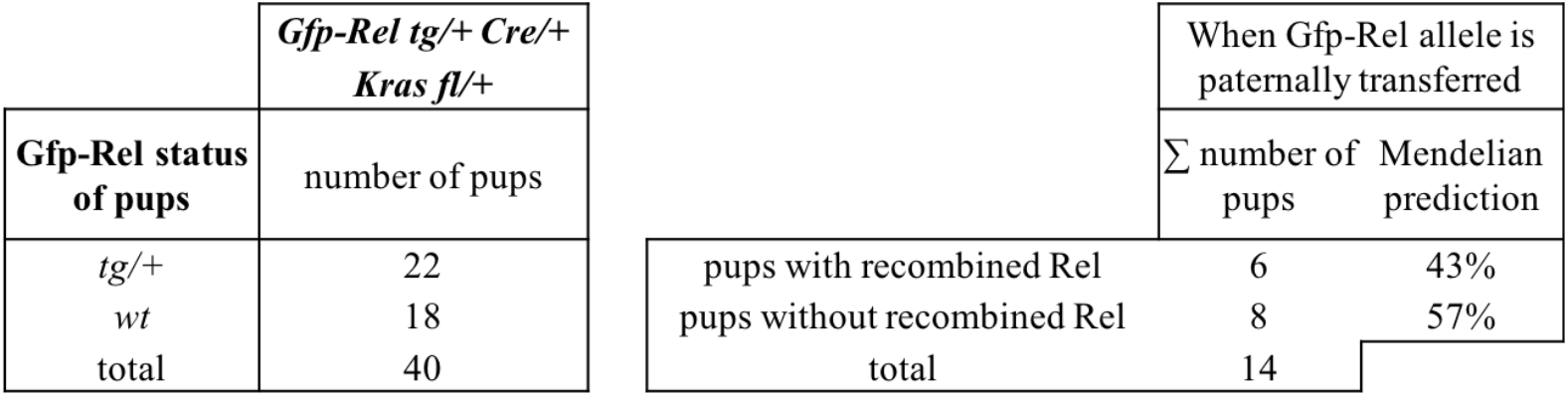
Number of mice obtained from GFP-Rel breeding

**Figure 3:**
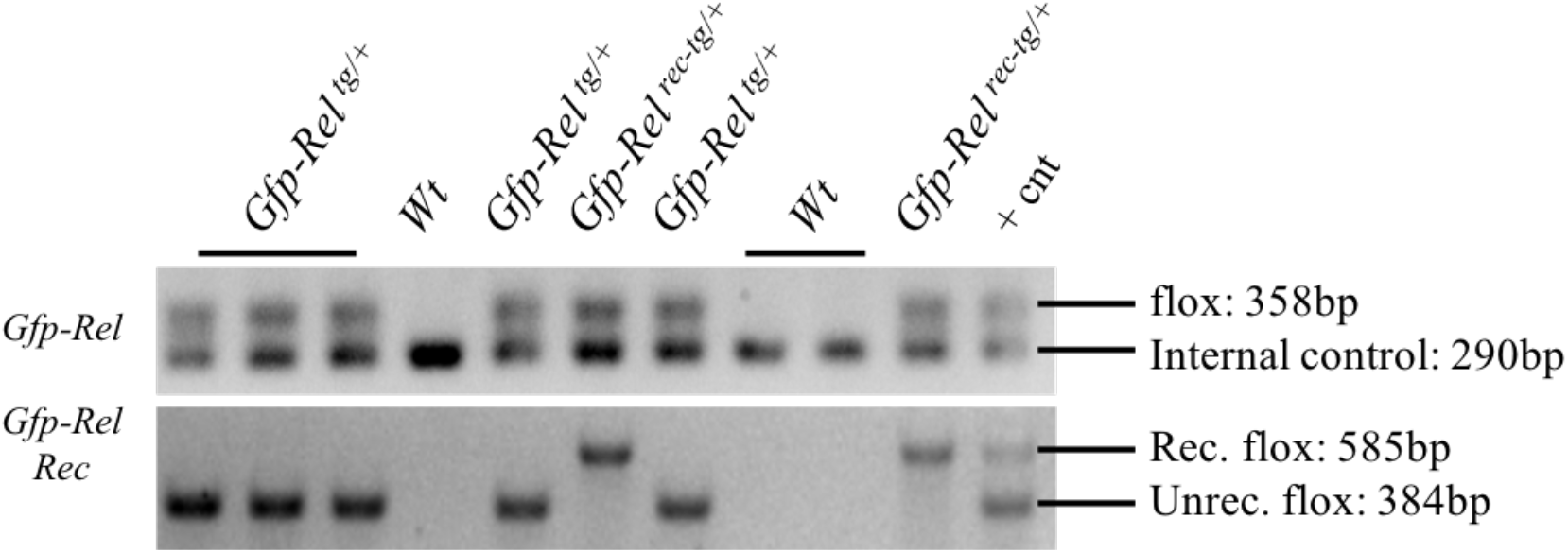
Representative PCR results obtained from GFP-Rel^*tg*^ mice breedings. A) Primer mix *Gfp-Rel* recognize the floxed allele along with an internal control. *Gfp-Rel Rec* primer mix gives band only if there is a *GFP-Rel* allele which can be recombined and unrecombined. **Rec**: recombined floxed allele. **Unrec**: unrecombined floxed allele. **rec-tg**: Recombined transgenic allele

## Discussion

In this work, we provided evidence that paternal transmission of *Cre* in *Ptf1a^tm1(cre)Hnak^* mouse model can cause germline recombination in the progeny. For the germline recombination, *Cre* transmission to progeny was not necessary. The germline recombination frequency changed depending on the target floxed allele. Neither *Kras* nor *Trp53* showed germline recombination while *RelB, Rel* and *GFP-Rel* showed in their respective mouse models with varying frequencies. No *Kras* recombination in the germline was observed. We postulate two possible reasons for this: recombination happens in the father germ cells before the fertilization occurs and the recombination efficiency of *Kras* is very low. Unlike other alleles, we didn’t observe any *Kras* recombination in the testes of the *Kras^fl/+^; Ptf1a-Cre/+* mice. Additionally, a difference in the *RelB* and *Kras* recombination efficiency can be observed in the pancreas PCRs (Figure 1D). While almost all of the floxed *RelB* allele was recombined in the *Ptf1a-Cre/+* pancreata, a big portion of *Kras* is still unrecombined.

Additionally, we observed recombination of the *Tp53* allele in the testes but not in the germline in *Trp53^tm1Brn^* model. A possible explanation is the acquired disadvantage for fertilization in the *Tp53* knock out sperms. Although *Tp53^-/-^* males are fertile, germ cell quality was reported to be regulated by *Tp53* [35–39]*. Trp53^tm2Tyj^* mouse model (LSL-Trp53^R172H^) has been highly used in pancreatic cancer research, especially in therapeutics [40]. Therefore, it is important to analyze whether or not it is recombined, as it can influence the drug response.

Depending on the gene, homozygous germline recombination might lead to embryonic lethality. However, many essential genes might still persist recombined heterozygously in the whole body which may or may not show any impact phenotypically. Especially in preclinical studies focusing on pancreatic pathology such as cancer and pancreatitis, a gross crosstalk is present between pancreatic and other cells (immune cells, neurons, blood vessels, stellate cells). Therefore, even if the heterozygous recombination of a protein is not observed phenotypically under normal circumstances, it may still impact the phenotype under pathological conditions.

Another highly used *Ptf1a*-driven mouse line is *Ptf1a^tm1.1(Cre)Cvw^* in which the entire *Ptf1a*-coding sequence was replaced with *Cre* [16], whereas in *Ptf1a^tm1(cre)Hnak^* Exon 1 was replaced [13]. In both of the mouse lines transcriptional regulatory elements before the coding sequence were retained. While we tested paternal recombination transmission only in *Ptf1a^tm1(cre)Hnak^*, we anticipate such an event to occur also in *Ptf1a^tm1.1(Cre)Cvw^*. In support of this, in JAX, paternal germline recombination transmission was already reported for an inducible *Ptf1a^tm2(cre/ESR1)Cvw^* model in which Exon-1 and 2 were replaced by *Cre-ERTM* [41].

Overall, we propose a necessity of genotyping for wt, floxed, and recombined alleles in all mice during regular mouse maintenance in all *Ptf1a-Cre* driven mouse lines. Accordingly, additional information regarding the genotyping protocols of the mouse models should be asked and provided by journals for good scientific practice. Unfortunately, for many of the mouse models there are no available primer sequences for the detection of recombined allele. Likewise, the maps and sequences of the targeted constructs are also in most of the cases not available for the individual design of primers. With this observation, we alert scientists, journals and reviewers to take immediate precautions for future studies.

## Materials and methods

### Mouse models

The mouse models used are *Ptf1a^tm1(cre)Hnak^* [13]; *Kras^tm4Tyj^* [42]; *Relb^tm1Fwei^* [43]; *Rel^tm1Ukl^* [44]; *GFP-Rel* (manuscript in revision, Marc Schmidt Supprian); *Trp53^tm1Brn^* [45]. Mice were kept in an animal room at room temperature (20-22°C) with light:dark cycle of 12:12 hours (light period: 06:00 am–06:00 pm) in groups of 2–4 animals in type III cages with bedding and nesting material. All animals were provided ad libitum with the standard food (No. 1324 – 10 mm pellets, Altromin, Lage, Germany) and water. All animals housed under specific pathogen-free conditions in accordance with the European Directive 2010/63/EU. All animal experiments were approved and conducted in accordance with the federal German guidelines for ethical animal treatment (Regierung von Oberbayern).

### Sample lysis and PCR

Collected samples are lysed in 200ul DirectPCR Lysis Reagent (Viagen) supplemented with 10ul PCR grade Proteinase K (Sigma) at 55°C. Once the samples are lysed, proteinase K is inactivated at 85°C for 1 hour. For PCR 2ul from the samples are directly used as template. For all of the PCRs, GoTaq® Green Master Mix (Promega) is used. For each reaction, 7.5ul 2X GoTaq master mix, 0.6ul primer mix (each 10uM) and 4.9ul nuclease free water are mixed. The cycling protocol is 95°C 1 min, (95°C 30 sec, 58°C 30 sec, 72°C 1.5 min) × 40 cycles, 72°C 10 min.

### Sample lysis and western blot

Tissues are homogenized and sonicated in NP-40 RIPA buffer (150mM NaCl, 1% NP-40, 0.5% Sodium deoxycholate, 0.1% SDS, 25mM Tris pH 7.4) with fresh protease and phosphatase inhibitors (Serva). 20-30μg per sample is run in 10% SDS-PAGE gel and transferred to 0.45μM PVDF membrane, blocked in 5% milk 1 hour RT and incubated overnight with primary antibodies diluted in milk. Next day they are incubated with HRP conjugated anti-rabbit secondary antibody for 1 hour at room temperature. Primary antibodies and their concentrations are as given: RelB sc-226 (C-19) Santa Cruz Biotechnology 1:500, GFP (ab183734) 1:1000 Abcam, c-Rel 12707 (D4Y6M) Cell Signaling Biotechnology 1:1000.

### Primer sequences

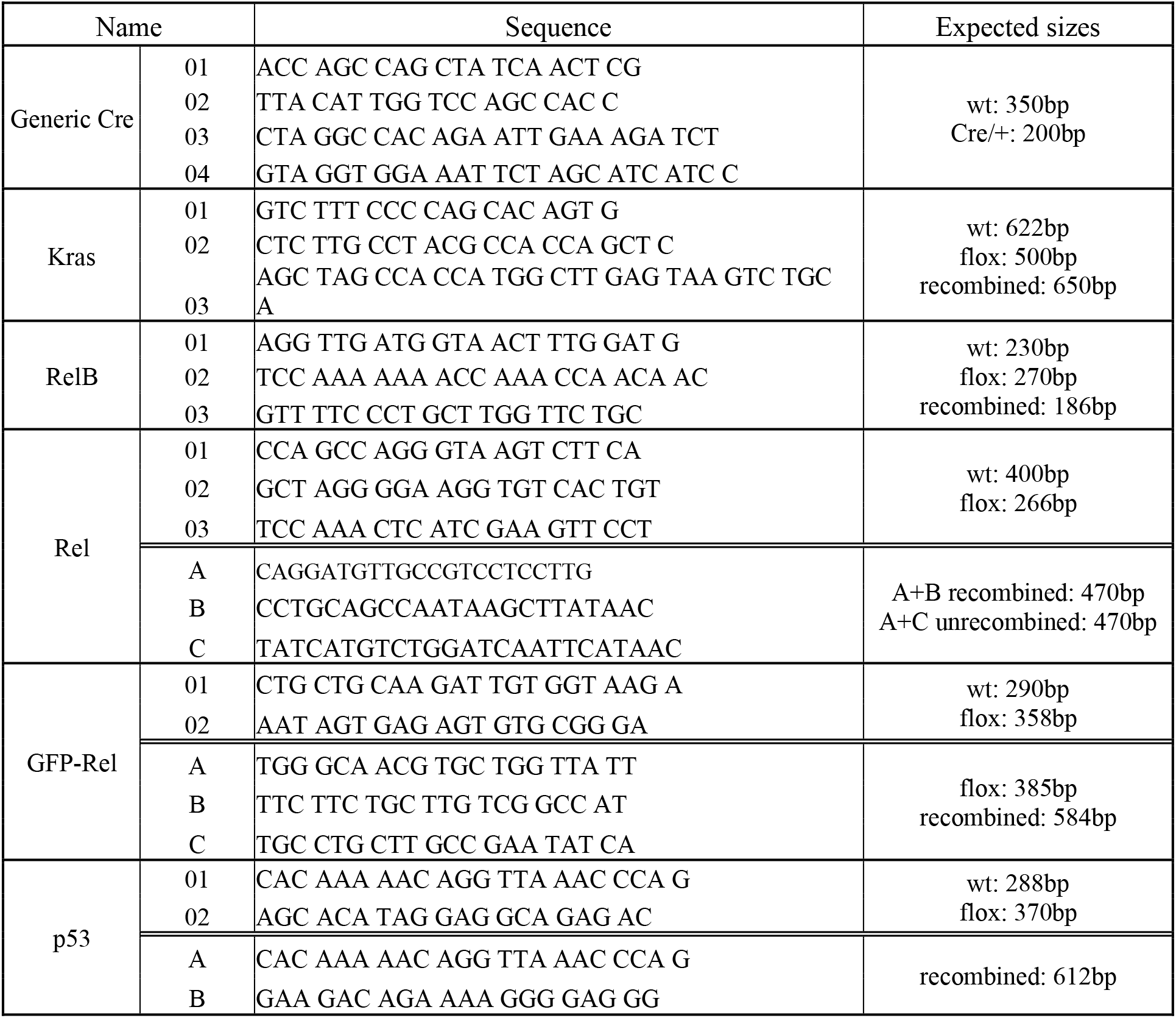

## Supplementary Figures

**Supplementary Figure 1:**
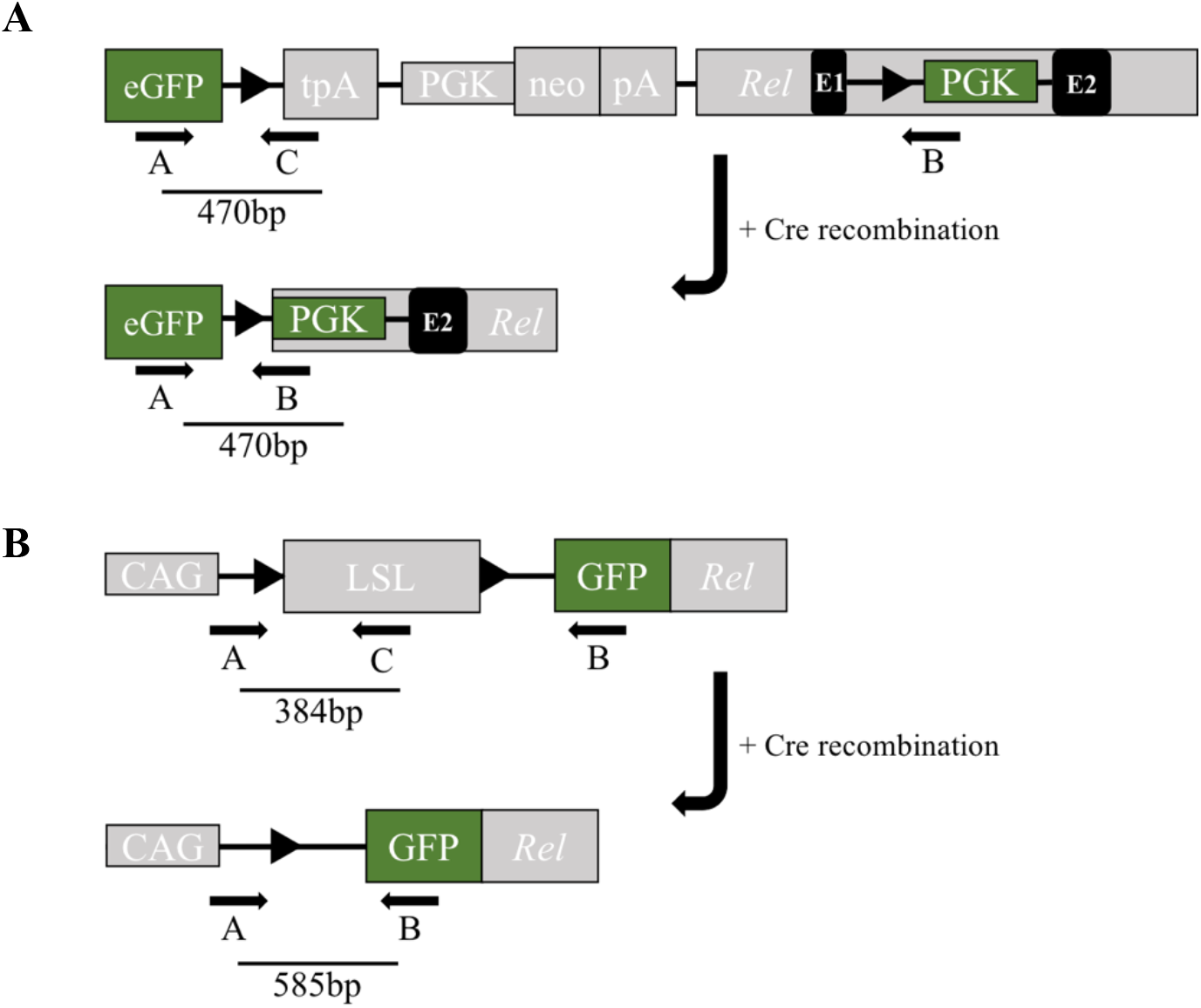
Schematics of the mouse lines with or without Cre recombination. A) A simplified map for Rel^tm1Ukl^ model. Exon1 of *Rel* is removed upon Cre mediated recombination, which brings together eGFP and PGK. Primer pair A+C recognizes the unrecombined allele with a size of 470bp. The distance between A-B pairs is too long to be amplified in unrecombined status. If there is recombination, A+B pair gives a band also at 470 bp. Primers sequences are kindly provided by Ulf Klein upon personal communication. B) A simplified map for GFP-Rel model. Upon Cre mediated recombination, constitutively active CAG promoter meets with N’ GFP-tagged *Rel*. As in Rel^tm1Ukl^ model, pair A+C recognizes unrecombined, and A+B recognizes recombined alleles with the given sizes. Primer sequences are designed based on the sequence map which is kindly provided by Marc Schmidt-Supprian.

## References

1. Sternberg, N.; Hamilton, D. Bacteriophage P1 site-specific recombination. I. Recombination between loxP sites. J. Mol. Biol. 1981, 150, 467–486.

2. Bohin, N.; Carlson, E.A.; Samuelson, L.C. Genome Toxicity and Impaired Stem Cell Function after Conditional Activation of CreERT2 in the Intestine. Stem Cell Reports 2018, 11, 1337–1346.

3. Janbandhu, V.C.; Moik, D.; Fässler, R. Cre recombinase induces DNA damage and tetraploidy in the absence of loxP sites. Cell Cycle 2014, 13, 462–470.

4. Gangoda, L.; Doerflinger, M.; Lee, Y.Y.; Rahimi, A.; Etemadi, N.; Chau, D.; Milla, L.; O’Connor, L.; Puthalakath, H. Cre transgene results in global attenuation of the cAMP/PKA pathway. Cell Death Dis 2012, 3, e365.

5. Pfeifer, A.; Brandon, E.P.; Kootstra, N.; Gage, F.H.; Verma, I.M. Delivery of the Cre recombinase by a self-deleting lentiviral vector: efficient gene targeting in vivo. Proc. Natl. Acad. Sci. U.S.A. 2001, 98, 11450–11455.

6. Loonstra, A.; Vooijs, M.; Beverloo, H.B.; Allak, B.A.; van Drunen, E.; Kanaar, R.; Berns, A.; Jonkers, J. Growth inhibition and DNA damage induced by Cre recombinase in mammalian cells. Proc. Natl. Acad. Sci. U.S.A. 2001, 98, 9209–9214.

7. Bersell, K.; Choudhury, S.; Mollova, M.; Polizzotti, B.D.; Ganapathy, B.; Walsh, S.; Wadugu, B.; Arab, S.; Kühn, B. Moderate and high amounts of tamoxifen in αMHC-MerCreMer mice induce a DNA damage response, leading to heart failure and death. Dis Model Mech 2013, 6, 1459–1469.

8. Forni, P.E.; Scuoppo, C.; Imayoshi, I.; Taulli, R.; Dastrù, W.; Sala, V.; Betz, U.A.K.; Muzzi, P.; Martinuzzi, D.; Vercelli, A.E.; et al. High levels of Cre expression in neuronal progenitors cause defects in brain development leading to microencephaly and hydrocephaly. J. Neurosci. 2006, 26, 9593–9602.

9. Li, Y.; Choi, P.S.; Casey, S.C.; Felsher, D.W. Activation of Cre Recombinase Alone Can Induce Complete Tumor Regression. PLoS One 2014, 9.

10. Schmidt, E.E.; Taylor, D.S.; Prigge, J.R.; Barnett, S.; Capecchi, M.R. Illegitimate Cre-dependent chromosome rearrangements in transgenic mouse spermatids. PNAS 2000, 97, 13702–13707.

11. Buerger, A.; Rozhitskaya, O.; Sherwood, M.C.; Dorfman, A.L.; Bisping, E.; Abel, E.D.; Pu, W.T.; Izumo, S.; Jay, P.Y. Dilated cardiomyopathy resulting from high-level myocardial expression of Cre-recombinase. J. Card. Fail. 2006, 12, 392–398.

12. McLellan, M.A.; Rosenthal, N.A.; Pinto, A.R. Cre-loxP-Mediated Recombination: General Principles and Experimental Considerations. Curr Protoc Mouse Biol 2017, 7, 1–12.

13. Nakhai, H.; Sel, S.; Favor, J.; Mendoza-Torres, L.; Paulsen, F.; Duncker, G.I.W.; Schmid, R.M. Ptf1a is essential for the differentiation of GABAergic and glycinergic amacrine cells and horizontal cells in the mouse retina. Development 2007, 134, 1151–1160.

14. Krapp, A.; Knöfler, M.; Ledermann, B.; Bürki, K.; Berney, C.; Zoerkler, N.; Hagenbüchle, O.; Wellauer, P.K. The bHLH protein PTF1-p48 is essential for the formation of the exocrine and the correct spatial organization of the endocrine pancreas. Genes Dev. 1998, 12, 3752–3763.

15. Rose, S.D.; Swift, G.H.; Peyton, M.J.; Hammer, R.E.; MacDonald, R.J. The role of PTF1-P48 in pancreatic acinar gene expression. J. Biol. Chem. 2001, 276, 44018–44026.

16. Kawaguchi, Y.; Cooper, B.; Gannon, M.; Ray, M.; MacDonald, R.J.; Wright, C.V.E. The role of the transcriptional regulator Ptf1a in converting intestinal to pancreatic progenitors. Nat. Genet. 2002, 32, 128–134.

17. Masui, T.; Long, Q.; Beres, T.M.; Magnuson, M.A.; MacDonald, R.J. Early pancreatic development requires the vertebrate Suppressor of Hairless (RBPJ) in the PTF1 bHLH complex. Genes Dev. 2007, 21, 2629–2643.

18. Masui, T.; Swift, G.H.; Deering, T.; Shen, C.; Coats, W.S.; Long, Q.; Elsässer, H.-P.; Magnuson, M.A.; MacDonald, R.J. Replacement of Rbpj with Rbpjl in the PTF1 complex controls the final maturation of pancreatic acinar cells. Gastroenterology 2010, 139, 270–280.

19. Holmstrom, S.R.; Deering, T.; Swift, G.H.; Poelwijk, F.J.; Mangelsdorf, D.J.; Kliewer, S.A.; MacDonald, R.J. LRH-1 and PTF1-L coregulate an exocrine pancreas-specific transcriptional network for digestive function. Genes Dev. 2011, 25, 1674–1679.

20. Obata, J.; Yano, M.; Mimura, H.; Goto, T.; Nakayama, R.; Mibu, Y.; Oka, C.; Kawaichi, M. p48 subunit of mouse PTF1 binds to RBP-Jkappa/CBF-1, the intracellular mediator of Notch signalling, and is expressed in the neural tube of early stage embryos. Genes Cells 2001, 6, 345–360.

21. Hoshino, M.; Nakamura, S.; Mori, K.; Kawauchi, T.; Terao, M.; Nishimura, Y.V.; Fukuda, A.; Fuse, T.; Matsuo, N.; Sone, M.; et al. Ptf1a, a bHLH transcriptional gene, defines GABAergic neuronal fates in cerebellum. Neuron 2005, 47, 201–213.

22. Sellick, G.S.; Barker, K.T.; Stolte-Dijkstra, I.; Fleischmann, C.; Coleman, R.J.; Garrett, C.; Gloyn, A.L.; Edghill, E.L.; Hattersley, A.T.; Wellauer, P.K.; et al. Mutations in PTF1A cause pancreatic and cerebellar agenesis. Nat. Genet. 2004, 36, 1301–1305.

23. Fujiyama, T.; Miyashita, S.; Tsuneoka, Y.; Kanemaru, K.; Kakizaki, M.; Kanno, S.; Ishikawa, Y.; Yamashita, M.; Owa, T.; Nagaoka, M.; et al. Forebrain Ptf1a Is Required for Sexual Differentiation of the Brain. Cell Rep 2018, 24, 79–94.

24. Kobayashi, Y.; Hensch, T.K. Germline recombination by conditional gene targeting with Parvalbumin-Cre lines. Front. Neural Circuits 2013, 7.

25. Liput, D.J. Cre-Recombinase Dependent Germline Deletion of a Conditional Allele in the Rgs9cre Mouse Line. Front. Neural Circuits 2018, 12, 68.

26. Zhang, J.; Dublin, P.; Griemsmann, S.; Klein, A.; Brehm, R.; Bedner, P.; Fleischmann, B.K.; Steinhäuser, C.; Theis, M. Germ-Line Recombination Activity of the Widely Used hGFAP-Cre and Nestin-Cre Transgenes. PLoS ONE 2013, 8, e82818.

27. Choi, C.-I.; Yoon, S.-P.; Choi, J.-M.; Kim, S.-S.; Lee, Y.-D.; Birnbaumer, L.; Suh-Kim, H. Simultaneous deletion of floxed genes mediated by CaMKIIα-Cre in the brain and in male germ cells: application to conditional and conventional disruption of Goα. Exp Mol Med 2014, 46, e93–e93.

28. Xie, C.; Zhu, F.; Wang, J.; Zhang, W.; Bellanti, J.A.; Li, B.; Brand, D.; Olsen, N.; Zheng, S.G. Off-Target Deletion of Conditional Dbc1 Allele in the Foxp3YFP-Cre Mouse Line under Specific Setting. Cells 2019, 8, 1309.

29. Luo, L.; Ambrozkiewicz, M.C.; Benseler, F.; Chen, C.; Dumontier, E.; Falkner, S.; Furlanis, E.; Gomez, A.M.; Hoshina, N.; Huang, W.-H.; et al. Optimizing Nervous System-Specific Gene Targeting with Cre Driver Lines: Prevalence of Germline Recombination and Influencing Factors. Neuron 2020, S0896627320300088.

30. Wu, D.; Huang, Q.; Orban, P.C.; Levings, M.K. Ectopic germline recombination activity of the widely used Foxp3-YFP-Cre mouse: a case report. Immunology 2020, 159, 231–241.

31. Spinelli, V.; Martin, C.; Dorchies, E.; Vallez, E.; Dehondt, H.; Trabelsi, M.-S.; Tailleux, A.; Caron, S.; Staels, B. Screening strategy to generate cell specific recombination: a case report with the RIP-Cre mice. Transgenic Res. 2015, 24, 803–812.

32. Rempe, D.; Vangeison, G.; Hamilton, J.; Li, Y.; Jepson, M.; Federoff, H.J. Synapsin I Cre transgene expression in male mice produces germline recombination in progeny. Genesis 2006, 44, 44–49.

33. Tsai, P.T.; Hull, C.; Chu, Y.; Greene-Colozzi, E.; Sadowski, A.R.; Leech, J.M.; Steinberg, J.; Crawley, J.N.; Regehr, W.G.; Sahin, M. Autistic-like behaviour and cerebellar dysfunction in Purkinje cell Tsc1 mutant mice. Nature 2012, 488, 647–651.

34. He, Y.; Sun, X.; Wang, L.; Mishina, Y.; Guan, J.-L.; Liu, F. Male germline recombination of a conditional allele by the widely used Dermo1-cre (Twist2-cre) transgene. Genesis 2017, 55.

35. Beumer, T.L.; Roepers-Gajadien, H.L.; Gademan, I.S.; van Buul, P.P.; Gil-Gomez, G.; Rutgers, D.H.; de Rooij, D.G. The role of the tumor suppressor p53 in spermatogenesis. Cell Death Differ. 1998, 5, 669–677.

36. Marty, M.S.; Singh, N.P.; Holsapple, M.P.; Gollapudi, B.B. Influence of p53 zygosity on select sperm parameters of the mouse. Mutat. Res. 1999, 427, 39–45.

37. Hasegawa, M.; Zhang, Y.; Niibe, H.; Terry, N.H.; Meistrich, M.L. Resistance of differentiating spermatogonia to radiation-induced apoptosis and loss in p53-deficient mice. Radiat. Res. 1998, 149, 263–270.

38. Hendry, J.H.; Adeeko, A.; Potten, C.S.; Morris, I.D. P53 deficiency produces fewer regenerating spermatogenic tubules after irradiation. Int. J. Radiat. Biol. 1996, 70, 677–682.

39. Yin, Y.; Stahl, B.C.; DeWolf, W.C.; Morgentaler, A. p53-mediated germ cell quality control in spermatogenesis. Dev. Biol. 1998, 204, 165–171.

40. Hingorani, S.R.; Wang, L.; Multani, A.S.; Combs, C.; Deramaudt, T.B.; Hruban, R.H.; Rustgi, A.K.; Chang, S.; Tuveson, D.A. Trp53R172H and KrasG12D cooperate to promote chromosomal instability and widely metastatic pancreatic ductal adenocarcinoma in mice. Cancer Cell 2005, 7, 469–483.

41. Kopinke, D.; Brailsford, M.; Pan, F.C.; Magnuson, M.A.; Wright, C.V.E.; Murtaugh, L.C. Ongoing Notch signaling maintains phenotypic fidelity in the adult exocrine pancreas. Dev. Biol. 2012, 362, 57–64.

42. Jackson, E.L.; Willis, N.; Mercer, K.; Bronson, R.T.; Crowley, D.; Montoya, R.; Jacks, T.; Tuveson, D.A. Analysis of lung tumor initiation and progression using conditional expression of oncogenic K-ras. Genes Dev. 2001, 15, 3243–3248.

43. Hamidi, T.; Algül, H.; Cano, C.E.; Sandi, M.J.; Molejon, M.I.; Riemann, M.; Calvo, E.L.; Lomberk, G.; Dagorn, J.-C.; Weih, F.; et al. Nuclear protein 1 promotes pancreatic cancer development and protects cells from stress by inhibiting apoptosis. J Clin Invest 2012, 122, 2092–2103.

44. Heise, N.; De Silva, N.S.; Silva, K.; Carette, A.; Simonetti, G.; Pasparakis, M.; Klein, U. Germinal center B cell maintenance and differentiation are controlled by distinct NF-κB transcription factor subunits. J. Exp. Med. 2014, 211, 2103–2118.

45. Marino, S.; Vooijs, M.; van Der Gulden, H.; Jonkers, J.; Berns, A. Induction of medulloblastomas in p53-null mutant mice by somatic inactivation of Rb in the external granular layer cells of the cerebellum. Genes Dev. 2000, 14, 994–1004.

